# Intact messenger RNA exists in human blood plasma and urine, and their purified macromolecular compartments

**DOI:** 10.1101/2024.11.30.626091

**Authors:** Jasper Verwilt, Kimberly Verniers, Sofie De Geyter, Sofie Roelandt, Cláudio Pinheiro, An Hendrix, Pieter Mestdagh, Jo Vandesompele

## Abstract

It is generally assumed that extracellular long RNA molecules in biofluids are fragmented. Few studies have indirectly hinted at the existence of possibly functional, intact long RNA transcripts. In search for such RNA molecules, we developed a long-read full transcript sequencing workflow for low-input and low-quality samples. We applied our method to human blood plasma, urine, and their isolated macromolecular compartments, in parallel with total RNA sequencing. This approach enabled us to find intact messenger RNA molecules in human biofluids and macromolecular compartments. We showed that the full-length transcriptome of human urine and blood plasma differs, but we also reveal intact messenger RNA molecules shared between biofluids. In addition, we show that these intact molecules are differentially distributed over fractionated macromolecular compartments. This study provides a foundation for future extracellular RNA studies to elucidate the human biofluid full-length transcriptome.

## Introduction

The complex cellular composition of ribonucleic acid (RNA), or transcriptome, controls the cell’s state and fate. The transcriptome’s intricacy is exemplified by a wide variety of RNA biotypes differing in length, structure, and function. For the human mind to comprehend its complexity, and for historic reasons, RNA is divided into short and long RNA. Short RNA, such as microRNA (miRNA), transfer RNA (tRNA), YRNA, and vault RNA (vRNA), mostly regulate the cellular inner-workings by associations with proteins, DNA, and other RNA molecules. Long RNA is an umbrella term for messenger RNA (mRNA), long non-coding RNA (lncRNA), and circular RNA (circRNA). While mRNA acts as a template for protein production, most lncRNA exerts a regulatory function as RNA itself. As a unique case, covalently-closed circRNA molecules ‘sponge’ miRNA^1^, sequester proteins^2^, or code for small proteins^3^.

As cells live and die, RNA molecules from the inside enter the extracellular space, actively or passively. In that way, the RNA transcripts enter various human biofluids and are subjected to harsh, typically RNA-unfriendly conditions^4^. Unexpectedly, some RNA species remain remarkably stable in biofluids^5^. This curiosity is explained by the association and protection of RNA transcripts by macromolecular structures, such as extracellular vesicles (EVs), lipoprotein particles (LPPs), and proteins (to form ribonucleoproteins (RNPs))^6,7^. While most research has focused on short RNA in human blood plasma-borne EVs, a handful of studies have characterized the long RNA transcripts across the macromolecular spectrum of human biofluids^6,8,9^.

Although extracellular long RNA is mostly fragmented^9–12^, some studies provide clues for the existence of a small proportion of intact molecules^8,12^. If intact extracellular mRNA exists, circulation could transport them from donor to acceptor cells, where they could be translated into their encoded protein, as demonstrated by in vitro experiments^13–15^.

Almost all current evidence for full-length extracellular mRNA in biofluids is indirect as it originates from short-read RNA sequencing (RNA-seq) data that fails to sequence long RNA molecules in their entirety. In contrast, long-read RNA-seq technologies, such as Oxford Nanopore Technologies (ONT) sequencing, determine the sequence of single RNA or cDNA molecules end-to-end, providing an direct and integral view of their sequence and structure. While ONT sequencing may in principle provide evidence for RNA transcript fragmentation status, the RNA content of most human biofluids is too low for PCR-based cDNA ONT sequencing.

To address this issue, we developed a low-input ONT sequencing protocol to characterize single and complete RNA molecules in human biofluids and fractionated macromolecular complexes. We use our newly developed method together with short-read RNA-seq to establish the polyadenylated RNA transcriptome of human platelet-free plasma and urine. Combining our approach with size and density purification, we provide further direct evidence for intact mRNA and lncRNA in biofluid-derived RNA-carrying macromolecular structures.

## Results

### Development of a low-input long-read full transcript sequencing workflow

Platelet-free plasma (PFP) and urine contain extremely low concentrations of messenger RNA (about 13-16 pg/mL)^5^. Due to their high input requirements, current Oxford Nanopore Technologies (ONT) cDNA sequencing protocols are unsuitable. To accommodate this, we combined various approaches to develop a low-input, quantitative long-read RNA sequencing method for low-quality samples (Figure 1B). Depending on the sample type and the input volume, we used a different RNA extraction kit. During the RNA extraction, intact Sequin spike-in RNA molecules were added as a control to verify absence of extraction-induced RNA fragmentation (Supplemental Figure 1A). To the RNA elute intact ERCC spike-in RNA molecules were added for control for library preparation-induced fragmentation (Supplemental Figure 1B). To remove any contaminating gDNA^16^, DNase treatment was performed, the success of which can be evaluated through data analysis (Supplemental Figure 2A, Supplemental Figure 7A). Amplified libraries were prepared using the oligo(dT)-based SMART-seq with UMI kit (Takara), which is optimized for low RNA concentrations. Finally, barcoded ONT sequencing adapters were ligated to the libraries for long-read sequencing using the SQK-RBK114 kit. The libraries were sequenced on PromethION flow cells in short-fragment mode. For the data analysis, we developed a Nextflow and Docker-reliant pipeline that robustly quantifies and characterizes the end-to-end-sequenced, polyadenylated reads (see Methods). The following sections show how we applied our method to characterize the full-length transcriptome of human plasma, urine, and their extracellular macromolecules.

**Figure 1:**
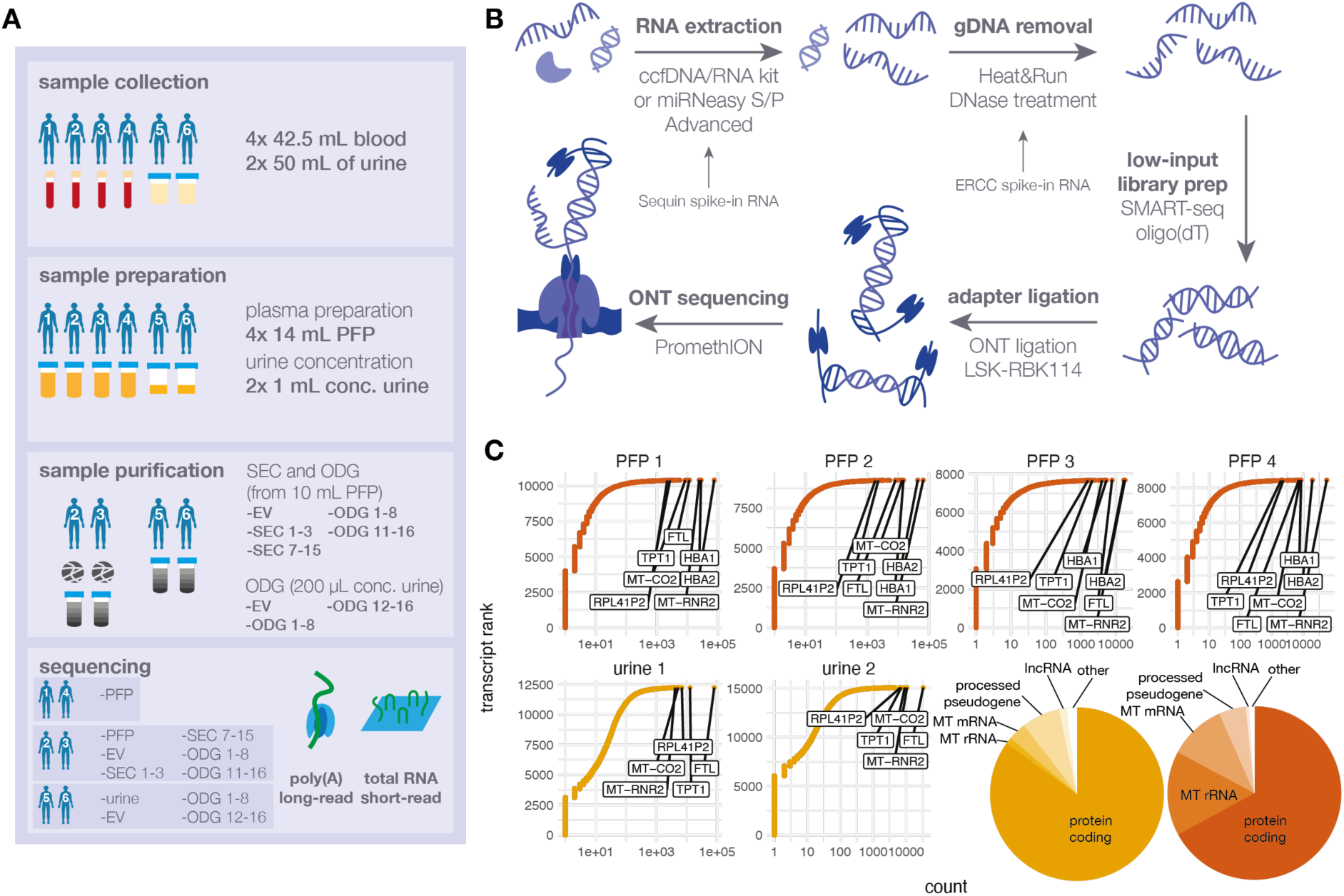
Experiment design, library preparation, and count inspection. (A) An overview of the experiment design of this study. SEC: size exclusion chromatography; ODG: OptiPrep density gradient; PFP: platelet-free plasma; conc. urine: concentrated urine. (B) Long-read sequencing workflow, from sample to sequencing. ccfDNA/RNA kit: QIAamp ccfDNA/RNA Kit (Qiagen); miRNeasy S/P: miRNeasy Serum/Plasma Advanced Kit (Qiagen); Heat&Run: Heat&Run gDNA removal kit (ArcticZymes); SMART-seq: SMART-seq mRNA LP (with UMI) (Takara Bio); ONT: Oxford Nanopore Technologies. (C) Count distributions for the transcripts detected in urine and PFP. The x-axis shows the raw counts of the transcript isoforms, the y-axis shows the ranking of the isoforms according to the number of raw counts (the x-axis). Each dot is a transcript isoform and the highest abundant transcripts in a given sample are indicated in that particular and the other samples. Two piecharts for each of the samples (urine in yellow, PFP in red-orange) show the five most abundant biotypes, the remaining biotypes are grouped under ‘other’.

### Long-read sequencing allows for in-depth characterization of polyadenylated RNA in blood plasma and urine

RNA in human biofluids is generally thought to be mainly fragmented, and existing evidence for a small portion of intact RNA remains indirect. We used our newly developed long-read full transcript sequencing approach in combination with short-read total RNA sequencing to characterize and quantify the intact transcriptome of human platelet-free plasma (PFP) and urine (Figure 1). We first verified the absence of DNA contamination by characterizing the strandedness of the short-read sequencing data (Supplemental Figure 2A). While our primary effort focused on quantifying the number of intact molecules in these biofluids using the spiked RNA molecules, we did not feel comfortable due to a number of biases that may result from the differential composition of the spike-in RNA and extracellular RNA. The count distributions are similar (Supplemental Figure 2B-C). The long-read counts correlate well with the short-read counts for protein coding genes (Supplemental Figure 2D). Most detected molecules in long-read sequencing data of PFP correspond to mRNA (67.5%, s.d. 11.3%), MT-rRNA (15.4, s.d. 7.22%), MT-mRNA (10.4%, s.d. 3.72%), processed pseudogenes (5.25%, s.d. 1.77%), and lncRNA (0.62%, s.d. 0.12%), while urine samples contained a higher fraction of mRNA (85.5%, s.d. 4.14%), processed pseudogenes (7.23%, s.d. 1.48%), and lncRNA (1.44%, s.d. 0.06%), and less MT-rRNA (1.19%, s.d. 0.90%) and MT-mRNA (3.43%, s.d. 1.52%) (Figure 1C). We detected an average of 8542 isoforms (s.d. 1181) from 6301 genes (s.d. 679), and 10 046 (s.d. 275) isoforms from 7336 genes (s.d. 173) in human PFP and urine, respectively, of which a total of 8557 isoforms are detected in at least one of the samples in both sample types. In PFP, a large proportion of the reads map to *HBA1* and *HBA2* isoforms, while these transcripts are completely absent in the urine data (as expected) (Figure 1C). Some transcripts, such as *TPT1* and *FTL* are abundantly present in both sample types (Figure 1C). In conclusion, we show that long-read ONT sequencing can characterize the extracellular polyadenylated RNA from human biofluids, such as PFP and urine.

### Human blood plasma and urine hold full-length RNA transcripts

Singe molecule long-read sequencing, in principle, allows to determine if and to what extent intact extracellular RNA exists in human plasma. We approximate the intactness of a transcript by counting the number of bases in the theoretical transcript matched by that observed in the read. In the urine and PFP libraries, ERCC reads are the most intact, followed by sequin and endogenous reads (Figure 2A). The coverage distributions for spike-in RNA are similar for the two biofluids. The spike-in RNA gives us an estimation of how well our method covers long transcripts. We find an inverse relationship between the fraction of the molecule that is covered and the theoretical length of the transcript (Supplemental Figure 3). For our integrity criteria, about 44.5% (s.d. 8.87%) and 34.3% (s.d. 5.90%) of all transcripts are intact in PFP and urine, respectively (Figure 2B). By far, most full-length reads originate from protein-coding genes (Figure 2C). In human PFP, we detected an average of 1032 (s.d. 157) unique, full-length isoforms per sample. In urine, we found an average of 816 (s.d. 3.54) unique, full-length isoforms per sample. In PFP, most full-length reads are assigned to hemoglobin (*HBA1, HBA2*), thymosin beta (*TMSB4X, TMSB10*), and ferritin (*FTL*) genes (Figure 2C-D). *TMSB4X, TMSB10,* and *FTL* are all also abundant and intact in human urine (Figure 2C-D). In summary, our approach provides direct evidence of intact polyadenylated RNA transcripts in human platelet-free blood plasma and urine.

**Figure 2:**
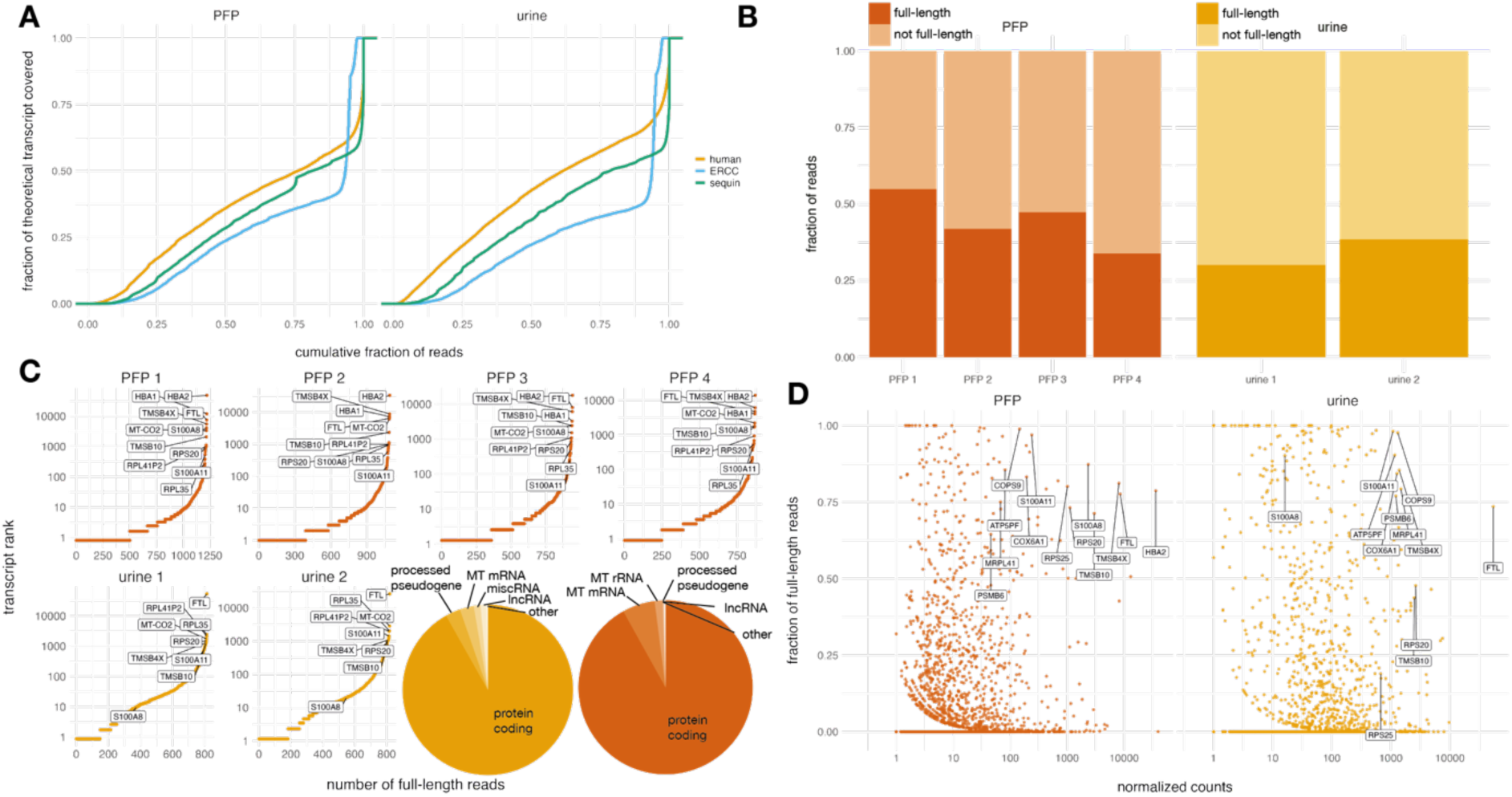
Exploration of intact polyadenylated RNA in human blood plasma and urine. (A) Length distribution of human, sequin spike-in, and ERCC spike-in reads. The x-axis shows the fraction of bases of the theoretical transcript matched by the reads. The y-axis shows the density distribution colored by read origin. (B) The distribution of full-length and fragmented transcripts in PFP and urine. For each donor, the y-axis shows the fraction of fully sequenced and uniquely mapping reads. (C) The number of full-length mRNA counts per sample. The x-axis shows the number of full-length reads for each transcript, the y-axis shows the ranking of that transcript (ranked by x-axis value). Each dot is a transcript isoform and the highest expressed transcripts in a given sample are indicated in that particular and the other samples. Two piecharts for each of the samples (urine in yellow, PFP in red-orange) show the five biotypes with the most intact reads, the remaining biotypes are grouped under ‘other’. (D) The average fraction of intact reads and DESeq2 normalized read counts for all isoforms. Each dot is an RNA isoform. The y-axis shows the fraction of reads that are full-length, averaged per isoform over all donors. The x-axis shows the normalized counts. Highly abundant isoform that have a minimal of 70% in any of the two sample types are indicated in both.

### Human blood plasma fractions display different full-length RNA repertoires

To better understand how intact RNA can survive a hostile environment, we separated the blood plasma into macromolecular fractions based on density and size. To this extent, we purified extracellular vesicles (EVs) from donors 3 and 4 using size exclusion chromatography (fractions 4-6) and density gradient separation (fractions 9-11) but also pooled fractions not corresponding to EVs as SEC 1-3, SEC 7-15, ODG 1-8, and ODG 12-16 (Figure 3A). The purification approach is identical to the one used by Vergauwen et al.^18^, and according to their characterization, SEC 1-3 (largest size) contains no lipoprotein particles (LPPs) but may contain larger EVs, such as apoptotic bodies; SEC 7-15 contains no EVs, but is enriched in LPPs (mainly HDL, and to a lesser degree VLDL, LDL, IDL) and abundant plasma proteins; ODG 1-8 (similar size, lower density compared to EVs) contains the remaining LPPs, and ODG 12-16 (similar size, higher density) contains high-density EVs and protein aggregates^18–20^. The abundance levels in these fractions do not correlate as well as in the original PFP samples, which is to be expected as the purification steps introduce additional variability and lower the RNA content (Pearson’s R between 0.84-0.97) (Supplemental Figure 4A).

**Figure 3:**
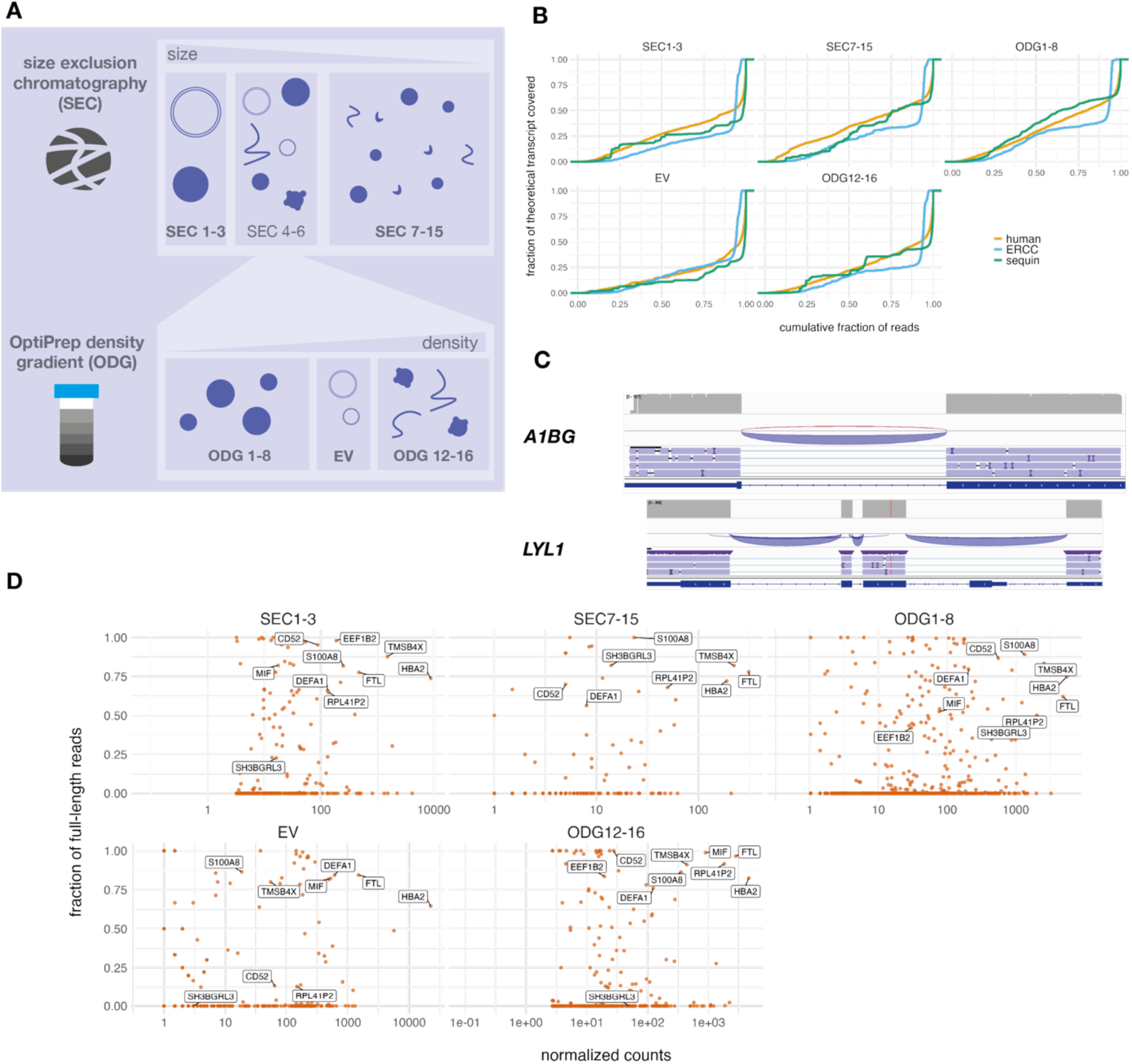
Full-length polyadenylated RNA sequencing in human blood plasma fractions. (A) An overview of the extracellular vesicle (EV) purification protocol and the composition of the remaining fractions. First, a size exclusion chromatography step (SEC) divides the macromolecules by size. The first three fractions (SEC 1-3) contain large EVs and chylomicrons, while the last nine fractions (SEC 7-15) contain small lipoprotein particles, such as high-density lipoprotein, and protein aggregates. The middle three SEC fractions (SEC 4-6) are pooled and further separated by density using a OptiPrep density gradient (ODG). The first eight fractions (ODG 1-8) contain most of the remaining lipoprotein particles, the last five ODG fractions (ODG 12-16) contain protein aggregates and other cellular debris. Fractions ODG 9-11 contain highly purified EVs. (B) Length distribution of human, sequin spike-in, and ERCC spike-in reads. The x-axis shows the fraction of bases of the theoretical transcript that is matched by the reads. The y-axis shows the density distribution colored by RNA source. (C) Integrative Genomics Viewer (IGV) screenshots of liver-specific gene *A1BG* and bone marrow and lymphoid tissue-enhanced gene *LYL1.* (D) The y-axis shows the average coverage for each isoform, the x-axis is the average normalized counts (averaged over donors). Each dot is an isoform and is colored by the fraction of intact transcripts. Gene names are displayed when an isoform is within the ten most abundant isoforms and has an intact fraction larger than 0.75.

The read coverage follows a similar pattern compared to the PFP and urine samples (Figure 3B). The fraction of intact reads varies among fractions: 47.8% (s.d. 4.26%) in SEC 1-3, 37.4% (s.d. 0.494%) in SEC 7-15, 30.8% (s.d. 5.38%) in ODG 1-8, 56.4.0% (s.d. 4.83%) in EVs, and 45.4% (s.d. 1.26%) in ODG 11-16 (Supplemental Figure 4B). Of these intact reads, the overwhelming majority originates from protein coding genes (Supplemental Figure 4C). Several intact transcripts are uniquely detected in one of the fraction types (Figure 3). *A1BG,* for example, a liver specific RNA (Human Protein Atlas, accessed 2023-10-07) is uniquely present as an intact transcript in ODG 1-8 (Figure 3C). Interestingly, while the protein product of *A1BG* is routinely found in human biofluids, such as plasma^21^, serum^22^, and synovial fluid_23_, the isoform detected in ODG 1-8 is a non-protein-coding, intron-retained transcript (A1BG-204). As another example, intact *LYL1* transcripts are only found in the EV fractions (Figure 3C). In tissues, *LYL1* is mainly expressed in bone marrow and lymphoid tissue (Human Protein Atlas, accessed 2023-11-07), and the extracellular abundance of its RNA transcripts has been used to predict prognosis in pancreatic ductal adenocarcinoma patients^24^. Most abundant mainly intact mRNA isoforms (average coverage larger than or equal to 75%), from genes such as *HBA2* and *FTL,* are present in all fractions (Figure 3D). In conclusion, our protocol can determine the full-length sequence of transcripts originating from lowly concentrated blood plasma-derived macromolecular fractions separated by sized and density and all fractions contain intact RNA molecules in varying degrees.

### Urinary macromolecules contain significantly less intact RNA

We extracted EVs from concentrated urine using a density gradient ultracentrifugation and pooled the remaining fractions: ODG 1-8 and ODG 12-16. ODG 1-8 is expected to contain mainly LPPs of varying densities and ODG 12-16 protein aggregates and fibers^20,28^ (Figure 4A). We performed long-read polyadenylated RNA sequencing on the EV and two pooled ODG fractions. Transcript counts within the same fraction type from different donors are strongly correlated (Supplemental Figure 5A). Fragmentation profiles of the human reads closely follow the spike-in RNA fragmentation profile (Figure 4B). Nevertheless, a lower fraction of the sequenced molecules are intact in urine compared to PFP (ODG 1-8: 19.1% (s.d. 0.928%), EV: 21.1% (s.d. 5.33%), and ODG 12-16: 10.7% (s.d. 6.06%)). Just like in PFP, highly abundant, mainly intact isoforms are shared between the various samples (Figure 4C). These results show the existence of intact mRNA transcripts in urine and how they are differentially distributed over its purified fractions.

**Figure 4:**
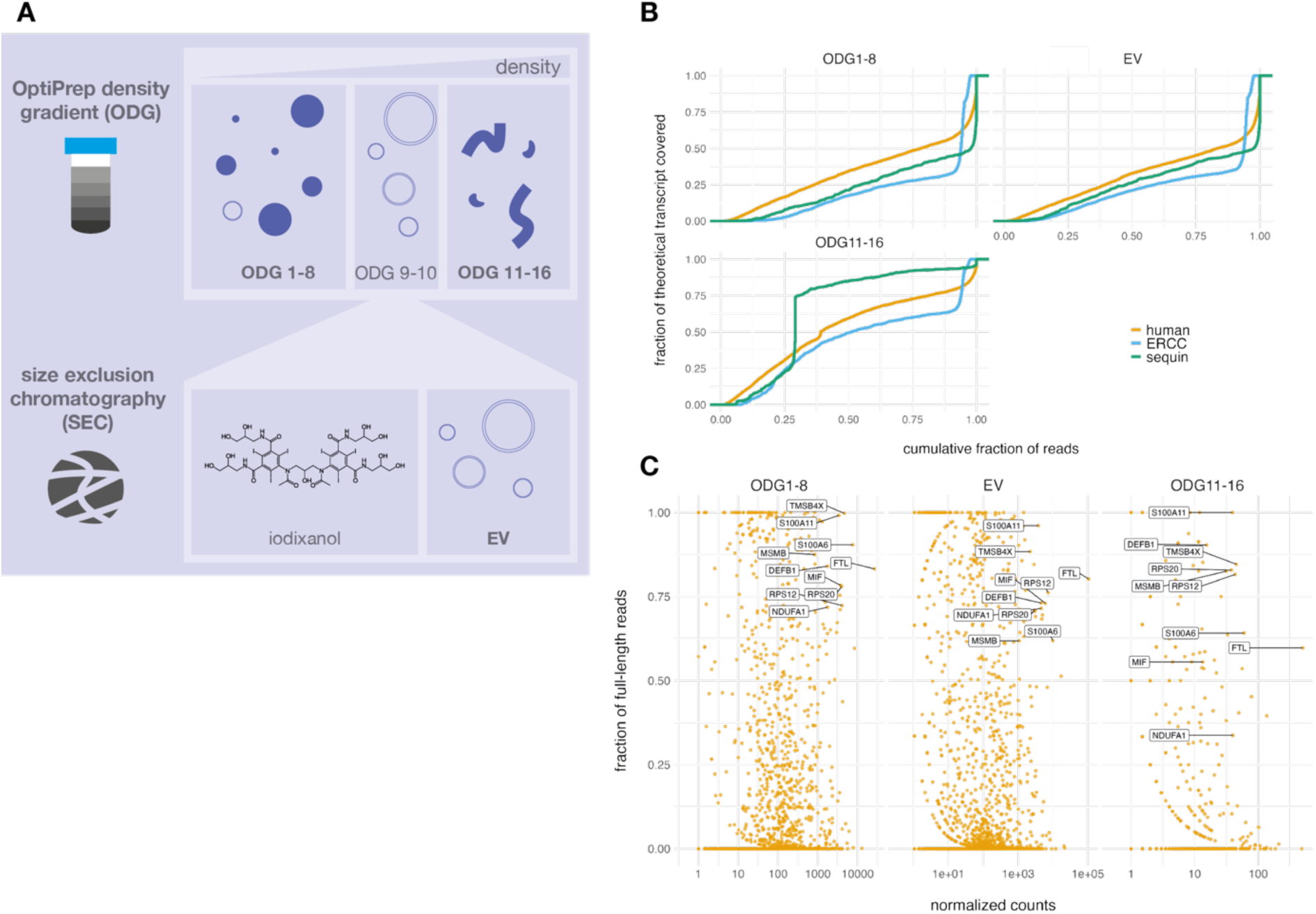
Full-length polyadenylated RNA sequencing in human urine and its density-separated fractions. (A) A summarized protocol for the isolation of EVs, along with the fractions that are pooled and sequenced. ODG 1-8 contains mainly lipoprotein particles, while ODG 11-16 contains mainly protein aggregates. (B) Integrity profiles for the different fractions. Density plots show the distribution of human, sequin spike-in, and ERCC spike-in reads in all sequenced fractions. The x-axis shows the fraction of bases of the theoretical transcript that is matched by the reads. The y-axis shows the density distribution colored by RNA source. (C) The y-axis shows the average coverage for each isoform, the x-axis is the average normalized counts (averaged over donors). Each dot is an isoform and is colored by the fraction of intact transcripts. Gene names are displayed when an isoform is within the ten most abundant isoforms and has an intact fraction larger than 0.70 in at least one of the samples.

### Urinary and blood plasma extracellular vesicles show transcriptome similarities

Since we purified EVs for both biofluids, we wondered to what extent the transcriptomes of these fractions corresponded with each other. We found 68 isoforms with at least one intact read in urine and PFP EVs and 58 and 1190 isoforms uniquely intact in PFP and urine, respectively (Figure 5A). While differences between both transcriptomes could be due to differences in separation and purification protocols or sequencing depth, similarities should be more robust. Of transcripts with intact reads in both samples, we find a group of genes for which the fraction of intact reads is relatively high in urinary and PFP EVs (Figure 5B). This group includes *MIF*, *FTL*, *TMSB4X*, *S100A8*, *RPS12*, and *RPS25.* Fragmentation profiles of other transcripts might not concur between two biofluids. While differences between both transcriptomes could be due to differences in separation and purification protocols or sequencing depth, stringent quality control of both purifications limits concern for the former issue. By focusing on the abundant transcripts, we remove large sequencing depth biases. The two purified fractions show differences and unique, intact transcripts (Figure 5C). In blood plasma-derived EVs, for example, we find a group of intact transcripts including two defensin alpha transcripts from *DEFA1* and *DEFA3*, which are enriched in Kuppfer cells (a type of neutrophil) (Human Protein Atlas, 14/10/’24). In urinary EVs, we find intact isoforms from, for example, *ATOX1* and *MT1X*, genes enriched in proximal tubular cells (Human Protein Atlas, 14/10/’24), which reside in the nephron. So, the EV populations in human blood plasma and urine are similar, but not identical and contain biofluid-dependent intact mRNA molecules.

**Figure 5:**
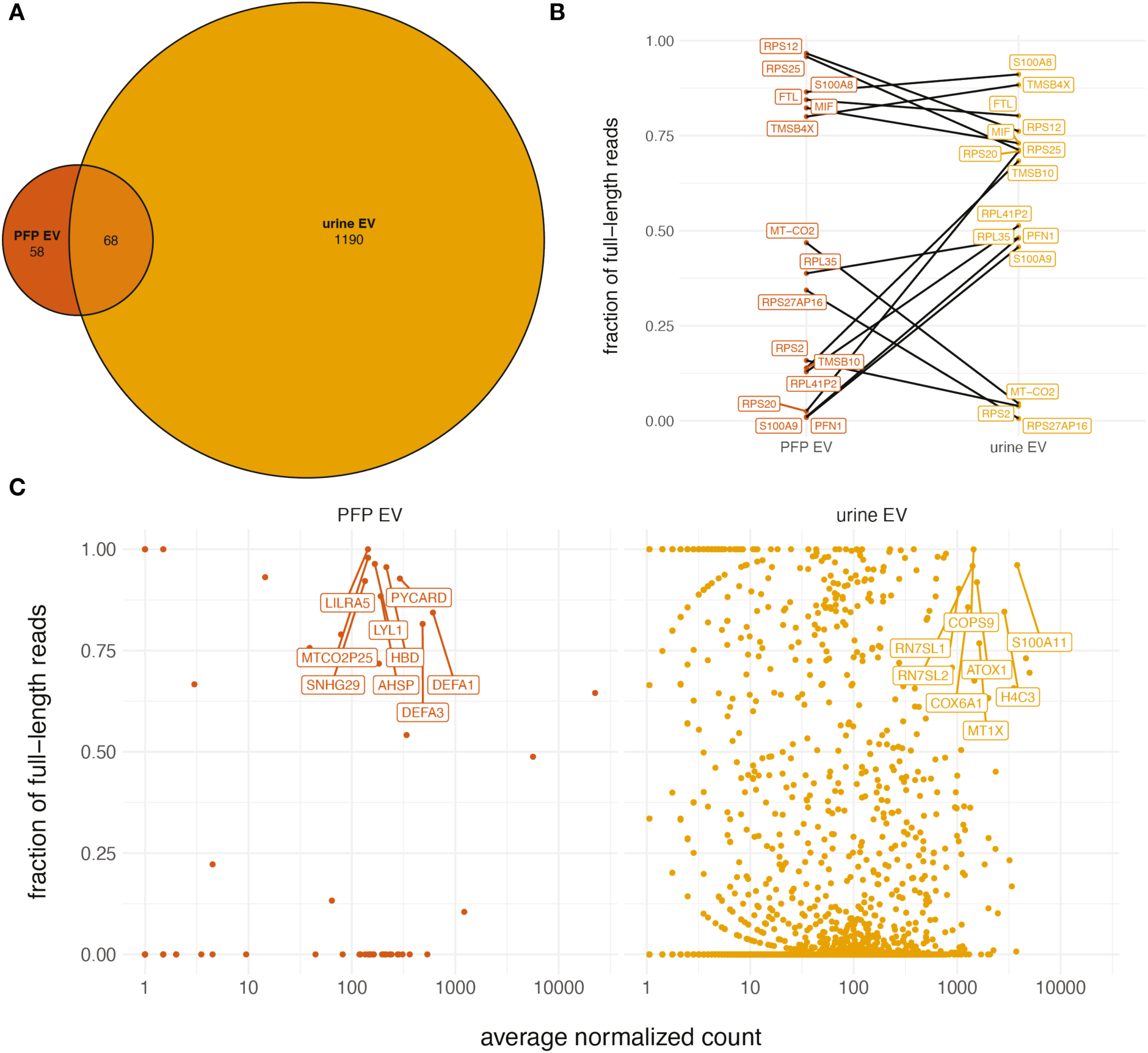
Comparison of PFP and urine extracellular vesicle-derived intact messenger RNA. (A) Overlap of isoforms with at least one intact read in both biological replicates for PFP EVs and urinary EVs. (B) Comparison of shared full-length isoforms between urinary and PFP EVs. The isoforms are filtered to have at least 10 raw counts in all biological replicates. The y-axis shows the-fraction of the reads that were full-length (calculated with the total number of reads over both biological replicates). Parent gene names of the isoforms are annotated and lines are drawn between isoforms. (C) Fragmentation and abundance profiles of unique transcripts for urinary and PFP EVs. Each dot is a transcript, the y-axis shows the average fraction of reads that are determined to be full-length. The x-axis shows the average normalized count over the two biological replicates (log10-transformed). Genes names are indicated for reads with a full-length fraction higher or equal to 0.75 and at least 100 and 1000 average normalized counts for PFP and for urine, respectively.

## Discussion

In this study, we developed a low-input long-read RNA-seq library preparation method and applied this method to human platelet-free blood plasma and urine. In addition, we separated macromolecules, such as EVs and LPP, of both biofluids using size and density-based separation methods and determined the intact RNA repertoire in each of them. The application of our technique provides the first direct evidence for the existence of intact full-length long RNA molecules in human biofluids.

While this study provides the first direct evidence for full-length RNA in human blood plasma and urine-derived EVs, previous research hinted at their existence. Older studies relied on short-read sequencing methods to characterize and identify intact extracellular long RNA. To add to the claim of full-length mature mRNA, researchers often perform an RT-qPCR to confirm the intactness of a set of observed mRNAs. Such research has led to the indirect identification of long RNA in extracellular vesicles (EVs)^8,29,30^. One study found the intact RNAs in EVs to have a maximum length of 1 kilobase^29^, although larger intact transcripts have been detected in microvesicles^12^, and only their fragmented counterparts are found in exosomes^12^. A recent study provided direct evidence for full-length, intact exRNA in 0.8 µm-filtered blood^31^, but it was unclear which macromolecular structures these RNAs were associated with. Another recent study used long-read sequencing to show the existence of intact mRNA molecules associated with cell-line-produced EVs^32^. About 60% of the reads mapped to mRNA isoforms, and 10% of EV transcripts were reported to be intact^32^. In contrast, we determined that mare than 90% of long reads in blood plasma EVs map to mRNA isoforms, with more than 50% of the transcripts being intact. Such differences are likely not biological but originate from differences in blood collection tube, time between draw and plasma preparation, EV isolation, RNA extraction, library preparation, and data analysis. For example, unlike our bioinformatic analysis, Padilla et al. did not perform any filtering for partially sequenced or internally primed reads.

Our results have various important implications. First, we provide a straightforward protocol for single-molecule long-read sequencing of low-input and low-quality RNA. The protocol is suitable for a wide variety of biofluids and sample types and opens these samples up to the advantages of ONT sequencing. Second, we showed the existence of extracellular intact mRNA and other biotypes in human blood plasma and urine. This adds to the discussion on extracellular RNA function. Researchers have shown how mRNA transcripts transported through EVs can reach acceptor cells, where they are translated into proteins^13–15,33^. It is still unclear to what extent intact extracellular mRNA can be translated by acceptor cells *in vivo*. Even if these transcripts can enter and be translated by other cells *in vivo*, it does not necessarily mean that they can elicit a biological response. In addition, enrichment of transcripts also does not implicate direct functionality. Specific loading of preferential transcripts is also in line with the hypothesis of it being cellular waste^34^. Third, we show that the fraction of intact RNA differs among biofluid-derived fractions. This observation implies that different macromolecular structures preferentially associate with intact long RNA (Figure 4D, for example). However, from our current results, it remains unclear whether these intact RNA molecules exist inside the vesicle-like structures or on the outside.

Our study has several limitations. First, we are currently unsure to what extent reverse transcription artifacts (or others) enter into the data analysis and are interpreted as fragmented reads^35^. For example, the strand-switching primer can bind the RNA internally and generate a shorter artifact. Another possibility is the generation of internally primed RNA when using oligo(dT) reverse transcription primers. Indeed, we note that in fractions containing more fragmented RNA, the spike-in RNA molecules follow a similar distribution (Figure 3B, for example). This could imply that a part of the fragmentation occurred during the RNA extraction or library preparation. We try to remove any internal priming artifacts by filtering the IsoQuant output for the ‘correct_polya_site’ flag. Second, although unique molecular identifiers (UMIs) are included in the library preparation, we currently do not perform deduplication. The UMI is incorporated during the template switching and is part of the SMART-seq kit. This kit, however, used six random bases as UMI, in contrast to the UMIs used by ONT, which are longer and more patterned. This makes the process of identifying the UMI within the read more complex. However, because the sequences are there, we will try to use them in future expansion of this research.

In conclusion, we provide a robust long RNA-seq library preparation workflow fit for low-quality sample sequencing. Human biofluids, their purified EVs, and other macromolecular carriers contain a remarkable fraction of intact polyadenylated RNA that can be explored to answer fundamental and clinical questions concerning the human extracellular transcriptome.

## Methods

### Spike-in RNA characterization

For quality control measures, we determined the concentration and the size distribution of the Sequin and ERCC spikes. We characterized a 1/10 dilution of both spikes using the TapeStation RNA HS kit (Agilent Technologies, CA, USA) and compared the true size distribution to the theoretical one (as calculated using the manufacturer’s molarity and sequence files).

## Biofluid collection and processing

### Blood draw and plasma preparation

For the blood draws and plasma preparation, we meticulously followed the European Committee For Standardization’s 17442 technical specification (CEN/TS 17442) for cfRNA analysis. We drew blood from four over-night fasting healthy donors. We drew eight ACD-A tubes (Beckson Dickinson, NJ, USA) from each donor. We selected ACD-A tubes due to their excellent performance in the exRNAQC study^43^ and for their low platelet activiation^44^. We inverted the tubes ten times immediately after the blood draw. For each donor, we randomized the tubes to exclude the order in which they were drawn as a confounding factor for further analysis. We started the procedure to prepare platelet-free plasma (PFP) according to the International Society on Thrombosis and Haemostasis (ISTH) protocol^45^ within 5 minutes of blood draw. First, we centrifuged the blood tubes for 15 minutes at 2500 x *g* and 20 °C, moved the supernatant to a 15 mL Falcon Tube. Second, we centrifuged again for 15 minutes at 2500 x *g* and 20 °C. Immediately after the last centrifugation step, we removed the supernatant PFP and instantly froze it in liquid nitrogen. We stored the frozen plasma at -80 °C. A maximum of 80 minutes elapsed from the start of plasma preparation to freezing.

### Urine collection

We performed the urine sample preparation previously described^46,47^. Briefly, we collected 50 mL of urine from 2 healthy volunteers and centrifuged it for 10 min at 1000 x g and 4 °C. The supernatant was concentrated using a Centricon plus-70 centrifugal filter with 10 kDa molecular weight cut-off (MWCO) (Merck Life Science, Germany) and adjusted to 1 mL with phosphate-buffered saline (PBS) (Thermo Fisher Scientific, MA, USA) (pH 7.2) Urine samples were immediately processed for EV preparation.

### Urine fractionation

EV preparation from urine samples was performed via a bottom up OptiPrep density gradient ultracentrifugation (DGUC) as previously described^46,47^. 800 µL of concentrated urine was used for EV extraction. The remaining 200 µL was stored at -80 °C. Solutions of 5%, 10% and 20% iodixanol were made by mixing appropriate amounts of homogenization buffer (0.25 M sucrose, 1 mM EDTA, 10 mM Tris-HCL (pH 7.4)) and iodixanol working solution. The working solution was prepared by combining a working solution buffer (0.25 M sucrose, 6 mM EDTA, 60 mM Tris-HCl, pH 7.4). The 800 µL urine sample was resuspended in 3.2 mL working solution, obtaining a 40% iodixanol suspension, and layered on the bottom of a 16.8 mL open top polyallomer tube (Beckman Coulter, CA, USA). A discontinuous bottom up DGUC was prepared by overlaying the urine suspension with 4 mL 20%, 4 mL 10% and 3.5 mL 5% iodixanol solutions, and 1 mL PBS (pH 7.2), respectively. The DGUC was centrifuged for 18 h at 100,000 x g and 4 °C (Beckman Coulter, CA, USA). Afterward, DGUC fractions of 1 mL were collected from the top of using the Biomek 4000 workstation where EV-enriched fractions 9 and 10 (corresponding to 1.09 – 1.11 g/mL densities) were further processed by size exclusion chromatography (SEC)^48^. Density fractions 1-8 and 12-16 were also stored. For SEC preparation, Sepharose CL-2B (cat. no. GE17-0140-01, GE Healthcare) was stored in ethanol and washed three times overnight with PBS (Thermo Fisher Scientific, MA, USA) (pH 7.2) before use. A 10 mL syringe (Romed Holland, Netherlands) was used in which a nylon net with 20 µm pore size (Merck Life Science, Germany) was placed at the bottom. The syringe was then filled up to 10 mL with Sepharose CL-2B using a circular motion. To avoid drying the Sepharose beads, PBS was added and immediately followed by loading of EV-enriched fractions and collection of 1 mL eluates. SEC fractions 4-7 were collected and concentrated to 100 µL using an Amicon Ultra-2 mL centrifugal filter with a 10 kDa cut-off (Merck Life Science, Germany). EV preparations were stored at -80 °C until further use.

### Blood plasma fractionation

EV preparation from blood plasma samples was performed via top down Optiprep DGUC as previously described^18,28,44^. Briefly, crude EV extracts from blood plasma were prepared via SEC columns (cfr. EV preparation from urine samples). For donor 2 and 3, we loaded 10 SEC columns with 1 mL of PFP each. We stored SEC fractions 1-3 and 7-15 for later use. Fractions 4-6 were collected, combined per donor, and concentrated to 3 mL using an Amicon Ultra-2 mL centrifugal filter with a 10 kDa cut-off (Merck Life Science, Germany). Three discontinuous DGUCs were prepared by layering 4 mL of 40%, 4 mL of 20%, 4 mL of 10%, and 3.5 mL of 5% iodixanol solutions on top of each other in a 16.8 mL open top polyallomer tube. One milliliter of crude EV extract was layered on top of each density gradient and centrifuged for 18h at 100,000 x g and 4 °C (Beckman Coulter, CA, USA). After centrifugation, gradient fractions of 1 mL were collected from top to bottom using the Biomek 4000 automated workstation^48^. Fractions 9 and 10, corresponding to a buoyant density of 1.09 – 1.11 g/ml, were collected and pooled. Fractions 1-8 and 11-16 were stored at -80 °C for later use. Lastly, to remove iodixanol from EV samples, a second SEC was included whereby eluates 4-7 were pooled, concentrated to 100 µL, and stored at -80 °C until further use.

## RNA extraction and DNase treatment

### Plasma RNA extraction

We extracted RNA from 4 mL of platelet-free plasma from each of the donors (four samples in total) using the QIAamp ccfDNA/RNA Kit (Qiagen, Germany) according to the manufacturer’s protocol. After addition of RPL buffer, we added 2 µL of a 1/2500 dilution of RNA Sequin Mix B (Garvan Institute of Medical Research, Australia) (further referred to as sequin spikes). We eluted the DNA/RNA with 16 µL of RNAse-free water. Next, we added 2 µL of a 1/25000 dilution of ERCC RNA Spike-In Mix (Thermo Fisher Scientific, MA, USA) (further referred to as ‘ERCC spikes’). To remove the co-eluted cfDNA, we used the Heat&Run gDNA removal kit (ArcticZymes, Norway) according to the manufacturer’s protocol.

### RNA extraction of urine and fractions

We extracted RNA from the urine, urinary EVs, ODG pool of fractions 1-8 (ODG 1-8), and ODG pool of fractions 11-16 (ODG 11-16). For each donor, we adjusted the volume of the urine (about 200 µl) and urinary EVs (about 100 µl) to 600 µL by adding 1X PBS. We then used the miRNeasy Serum/Plasma Advanced Kit (Qiagen, Germany) according to the manufacturer’s protocol to extract RNA from 600 µL of each of the four samples. After addition of RPL buffer, we added 2 µL of a 1/375000, 1/1875000, 1/2812500, 1/750000, and 1/6300000 dilution of RNA Sequin Mix B (Garvan Institute of Medical Research, Australia) to urine, urinary EVs, ODG 1-8 and ODG 11-16 samples, respectively. We eluted the DNA/RNA with 16 µL of RNAse-free water. Next, we added 2 µL of a 1/3750000, 1/7031250, 1/28125000, 1/7500000, and 1/18000000 dilution of ERCC RNA Spike-In Mix (Thermo Fisher Scientific, MA, USA) urine, urinary EVs, ODG 1-8 and ODG 11-16 RNA, respectively. To remove the co-eluted cfDNA, we used the Heat&Run gDNA removal kit (ArcticZymes, Norway) according to the manufacturer’s protocol.

### RNA extraction of plasma fractions

We extracted RNA from the EVs, pool of SEC fractions 1-3 (SEC 1-3), pool of SEC fractions 7-15 (SEC 7-15), ODG pool of fractions 1-8 (ODG 1-8), and ODG pool of fractions 11-16 (ODG 11-16). For each donor, we adjusted the volume of the EVs (about 300 µl) to 4 mL by adding 1X PBS. We then used the QIAamp ccfDNA/RNA Kit (Qiagen, Germany) according to the manufacturer’s protocol to extract RNA from 600 µL of each of the four samples. After addition of RPL buffer, we added 2 µL of a 1/6000, 1/5000, 1/133200, and 1/99900 dilution of RNA Sequin Mix B (Garvan Institute of Medical Research, Australia) to urine, urinary EVs, ODG 1-8 and ODG 11-16 samples, respectively. We eluted the DNA/RNA with 16 µL of RNAse-free water. Next, we added 2 µL of a 1/180000, 1/450000, 1/444000, and 1/333000 dilution of ERCC RNA Spike-In Mix (Thermo Fisher Scientific, MA, USA) urine, urinary EVs, ODG 1-8 and ODG 11-16 RNA, respectively. To remove the co-eluted cfDNA, we used the Heat&Run gDNA removal kit (ArcticZymes, Norway) according to the manufacturer’s protocol.

## Polyadenylated RNA long-read sequencing

### mRNA long-read library preparation

We used the SMART-seq mRNA kit (with UMIs) (Takara Bio Europe, France) to generate c‡DNA libraries for the plasma, urine, and fractions. We followed the protocol according to the manufacturer’s instructions, with a few modifications. First, we elongated the bead incubation period to 10 minutes on a HulaMixer (Thermo Fisher Scientific, MA, USA), and the bead elution to 5 minutes on a HulaMixer (Thermo Fisher Scientific, MA, USA) and 5 minutes at 37 °C on a ThermoCycler (heated lid off). Second, we used SPRI beads in 0.75X volume ratio instead of AMPure beads. Third, we performed a varying number of PCR cycles: plasma 14, plasma SEC 1-3 27, plasma SEC 7-15 27, plasma ODG 1-8 29, plasma ODG 12-16 29, plasma EV 27, urine 28, urine ODG 1-8 28, urine ODG 11-16 28, and urine EV 28. After the final cleanup, we characterized the samples using the TapeStation HS D5000 kit (Agilent Technologies, CA, USA). We then used the Native Barcoding Kit 24 v14 (Oxford Nanopore Technologies, UK) to prepare the cDNA libraries for sequencing. We made the same adjustments as in the SMART-seq mRNA kit concerning the bead cleanup steps and included a cleanup step after barcoding to avoid barcode crossover. The plasma was barcoded 1-4, the plasma fractions were barcoded 1-10, and the urine and fractions were barcoded 1-12.

### PromethION sequencing

We loaded 15.24 fmol for plasma, 17.88 fmol for the plasma fractions, and 23 fmol for the urine fractions on a PromethION flow cell. The sequencing was done in short-fragment mode and the data was stored as POD5 files. Sequencing reports are available at https://github.com/OncoRNALab/plasmaONT.

## Short-read sequencing

### Library preparation

Libraries for all samples were prepared and sequenced in one experiment. We used 4.5 µL of the DNase-treated and spiked RNA to start the SMARTer Stranded Total RNA-Seq Kit v3 - Pico Input Mammalian (Takara Bio, USA). The kit was used according to the user’s manual, with a few adjustments: (1) all RNA was fragmented for 2 minutes at 94 °C, and (2) 16 PCR cycles were used for all samples. Libraries were then quantified using the KAPA Library Quantification Kit (Roche, Switzerland) on a LightCycler 480 (Roche, Switzerland) and characterized using an HS Small Fragment Kit (Agilent Technologies, CA, USA) on 5300 Fragment Analyzer System (Agilent Technologies, CA, USA). Next, we prepared an equimolar pool of all samples and proceeded with sequencing.

### Illumina sequencing

We loaded the library at a concentration of 650 pM with 2% PhiX and run in 2x50 basepairs paired-end mode on a NextSeq 2000 (Illumina, CA, USA). The sequencing resulted in 516 million reads of which 91.31% had a Q-score of Q30 or higher. The sequencing report and a MultiQC summary are available at https://github.com/OncoRNALab/plasmaONT.

## Data analysis

### Long-read sequencing analysis

We analyzed the POD5 files using our custom Nextflow-Docker pipeline (available at https://github.com/OncoRNALab/plasmaONT). To prepare the data for this pipeline, we start by re-basecalling all reads using Dorado^49^ (v0.3.1) (on a GPU), followed by conversion to FASTQ files with Samtools^50^ (v1.16.1) and demultiplexing using Porechop^51^ (v0.2.4) (without adapter trimming). The demultiplexed FASTQ files were used as input in our NextFlow pipeline. After combining all FASTQ files per sample, the reads files are channeled to two different processes. First, we Porechop_ABI^52^ (v0.2.4) to detect the adapter sequences *ab initio*. The detected adapter sequences are stored and trimmed from the reads. Then, feed the stored adapter sequences and the untrimmed reads to Pychopper^53^ (v2.7.9), which filters the data for end-to-end sequenced reads. We then map the Porechop_ABI and Pychopper-trimmed reads (two different channels) to the genome using Minimap2^54^ (v2.26.0). Last, only the Porechop_ABI-trimmed reads are quantified using IsoQuant^56^ (v3.3.0). We only retain reads tagged by IsoQuant to map to a poly(A) site. To acknowledge small insertions and other mismatches in the data, we define a transcript ‘intact’ if the read covers more than 85% of the theoretical isoform sequence, and having IsoQuant tags tes_match/tes_match_precise, tss_match/tss_match_precise, and fsm/mono_exonic_match.

### Short-read sequencing analysis

We used an in-house developed pipeline for the short-read sequencing data analysis. The pipeline starts by trimming adapters from the reads using cutadapt^57^ (v3.5). Next, the UMI sequences are extracted from the first read using UMI-tools^58^ (v1.0.1) and removed using cutadapt^57^ (v3.5). Next, we use Kallisto^59^ (v0.46.1) to pseudo-align the reads and quantify the expressed isoforms on a transcripts per million (TPM) level. The complete pipeline is available at https://github.com/OncoRNALab/plasmaONT.

### Results generation

For further analysis, we used R (v4.1.0). An RMarkdown file shows how the data is handled and how the figures and tables are generated (available at https://github.com/OncoRNALab/plasmaONT).

## Data availability

All raw data will be made available as soon as possible. Sequencing reports, bioinformatic pipelines, code, figures, and tables available at https://github.com/OncoRNALab/plasmaONT.

## Acknowledgements

We would like to thank NXTGNT (Ghent, Belgium) for carrying out the PromethION sequencing.

## Author Contributions

Conceptualization: A.H., J.Va., J.Ve., and P.M.; data curation: J.Ve.; formal analysis: J.Ve.; funding acquisition: A.H., J.Va., and P.M.; investigation: C.P., J.Ve., K.V., S.D.G., and S.R. methodology: A.H., J.Va., J.Ve., and P.M.; project administration: J.Va., J.Ve., and P.M.; resources: A.H., J.Va., P.M., and S.R.; software: J.Ve., supervision: J.Va. and P.M.; validation: J.Ve.; visualization: J.Ve.; writing—original draft: C.P., J.Ve.; writing—review & editing: A.H., C.P., J.Va., J.Ve., K.V., P.M., S.D.G., and S.R.

## Competing interests

J.Ve. has received funds for accommodation and travel to provide invited research presentations at Nanopore Day Ghent 2022 and Nanopore Community Meeting 2022 in New York, NY, USA.

## Notes

### Competing Interest Statement

The authors have declared no competing interest.

