## Supplemental Figures for "Intact messenger RNA exists in human blood plasma and urine, and their purified macromolecular compartments"

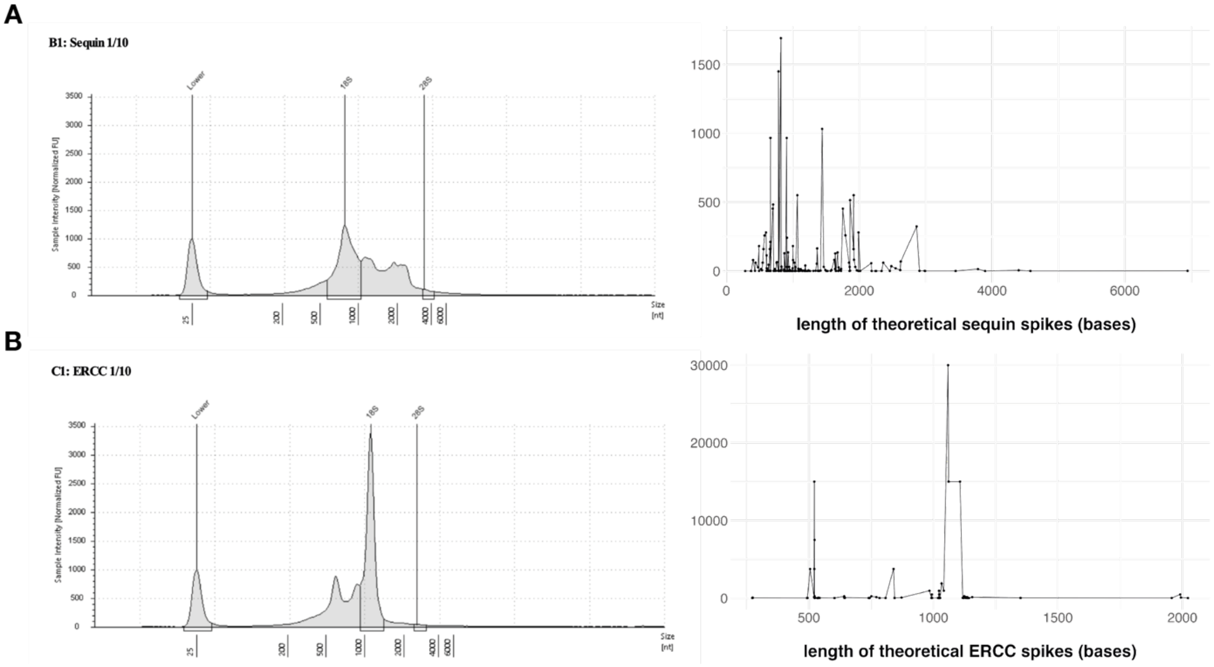

**Supplemental Figure 1: Spike fragmentation profiles.** The figures on the left show TapeStation fragmentation profiles of the 1/10 dilutions of the sequin (A) and ERCC (B) spikes used in the experiments. The theoretical fragment distributions based on the annotation are shown on the right.

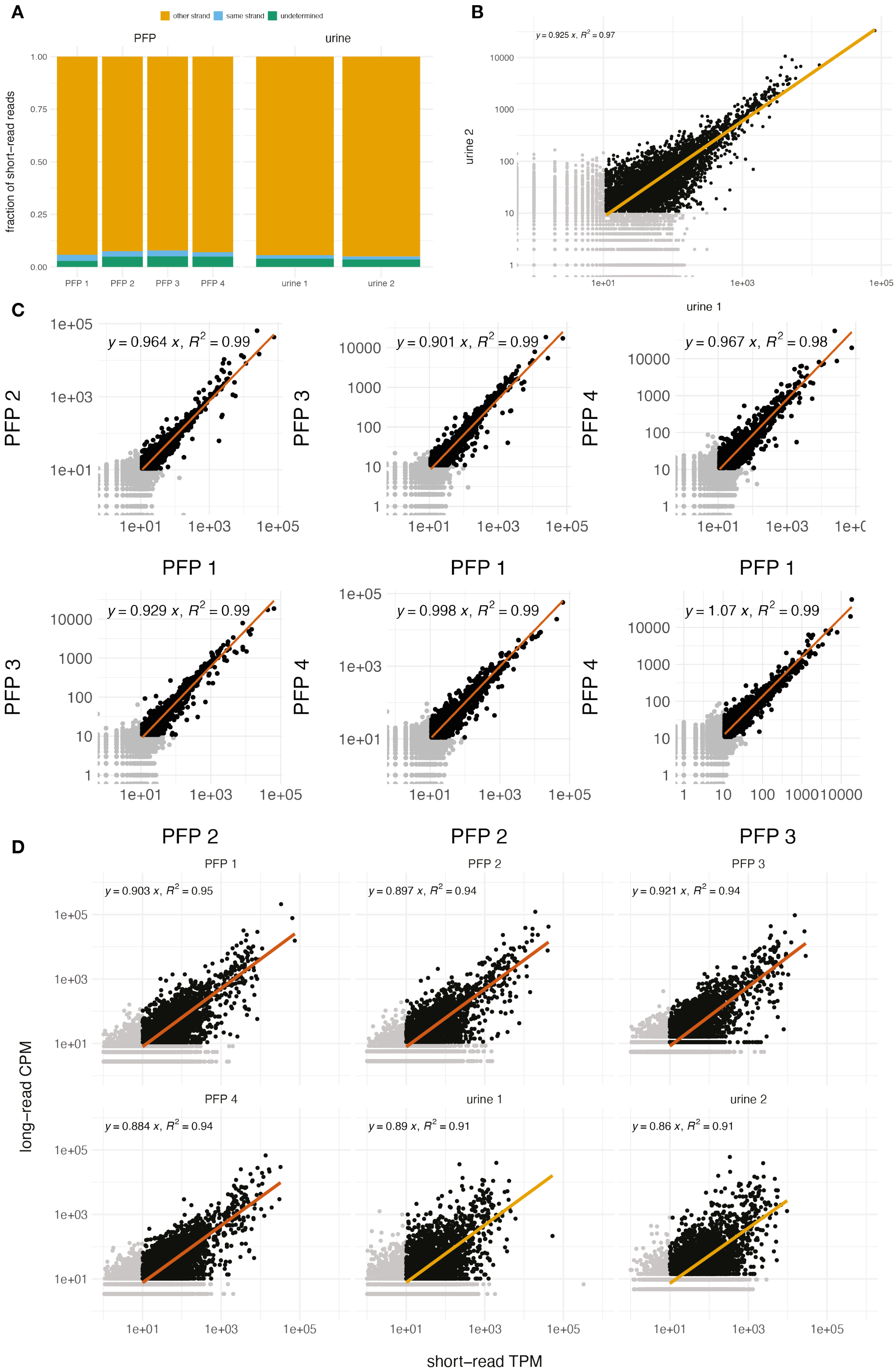
A
other strand
same strand
undetermined
PFP
urine
fraction of short−read reads
1.00
0.75
0.50
0.25
0.00
PFP 1
PFP 2
PFP 3
PFP 4
urine 1
urine 2
B
y = 0.925 x, R² = 0.97
urine 2
urine 1
C
y = 0.964 x, R² = 0.99
y = 0.901 x, R² = 0.99
y = 0.967 x, R² = 0.98
y = 0.929 x, R² = 0.99
y = 0.998 x, R² = 0.99
y = 1.07 x, R² = 0.99
PFP 2
PFP 3
PFP 4
PFP 1
D
y = 0.903 x, R² = 0.95
y = 0.897 x, R² = 0.94
y = 0.921 x, R² = 0.94
y = 0.884 x, R² = 0.94
y = 0.89 x, R² = 0.91
y = 0.86 x, R² = 0.91
PFP 1
PFP 2
PFP 3
PFP 4
urine 1
urine 2
long−read CPM
short−read TPM

**Supplemental Figure 2: Sequencing data exploration.** (A) For each donor, the y-axis shows the percentage of reads that are assigned as mapping to the same strand or a different strand from the coding strand. If this could not be determined, the reads are classified as 'undetermined'. (B) Comparison of raw long-read counts for both urine samples. Each dot is a transcript and are colored in gray if one the isoform had less than 10 counts in any of the compared samples, these are excluded from the regression analysis. Both axes are log10 transformed and the equation of the linear model (forced through 0) and $R^2$ are shown. The linear model is generated using the ggplot geom_smooth(method = "lm") layer. (C) Comparison of raw long-read counts for each of the PFP samples. Each dot is a transcript and are colored in gray if one the isoform had less than 10 counts in any of the compared samples. Both axes are log10 transformed and the equation of the linear model (forced through 0) and $R^2$ are shown. The linear model is generated using the ggplot geom_smooth(method = "lm") layer. (D) Comparison of CPM of the long-read data (raw counts / total) and TPM of the short-read data (length-normalized TPMs) filtered for each protein coding gene. Genes with TPM or CMP less than 10 are colored in gray and are not included in the regression analysis. Both axes are log10 transformed and the equation of the linear model (forced through 0) and $R^2$ are shown. The linear model is generated using the ggplot geom_smooth(method = "lm") layer.

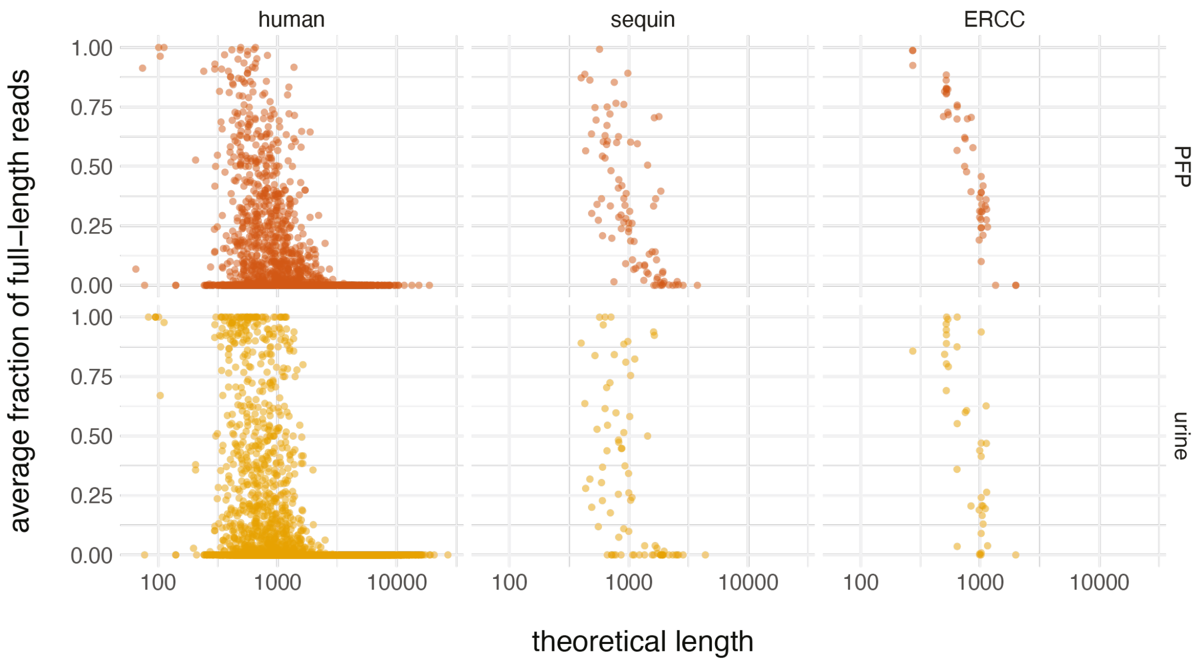


**Supplemental Figure 3: Relation of theoretical length of transcript and fraction of full-length reads** . For each transcript, and grouped by sample type and origin, the fraction of reads that are full-length is shown in relation to the theoretical isoform length. Only transcripts with at least 10 counts in any of the samples are included and their fractions of full-length reads are averaged.

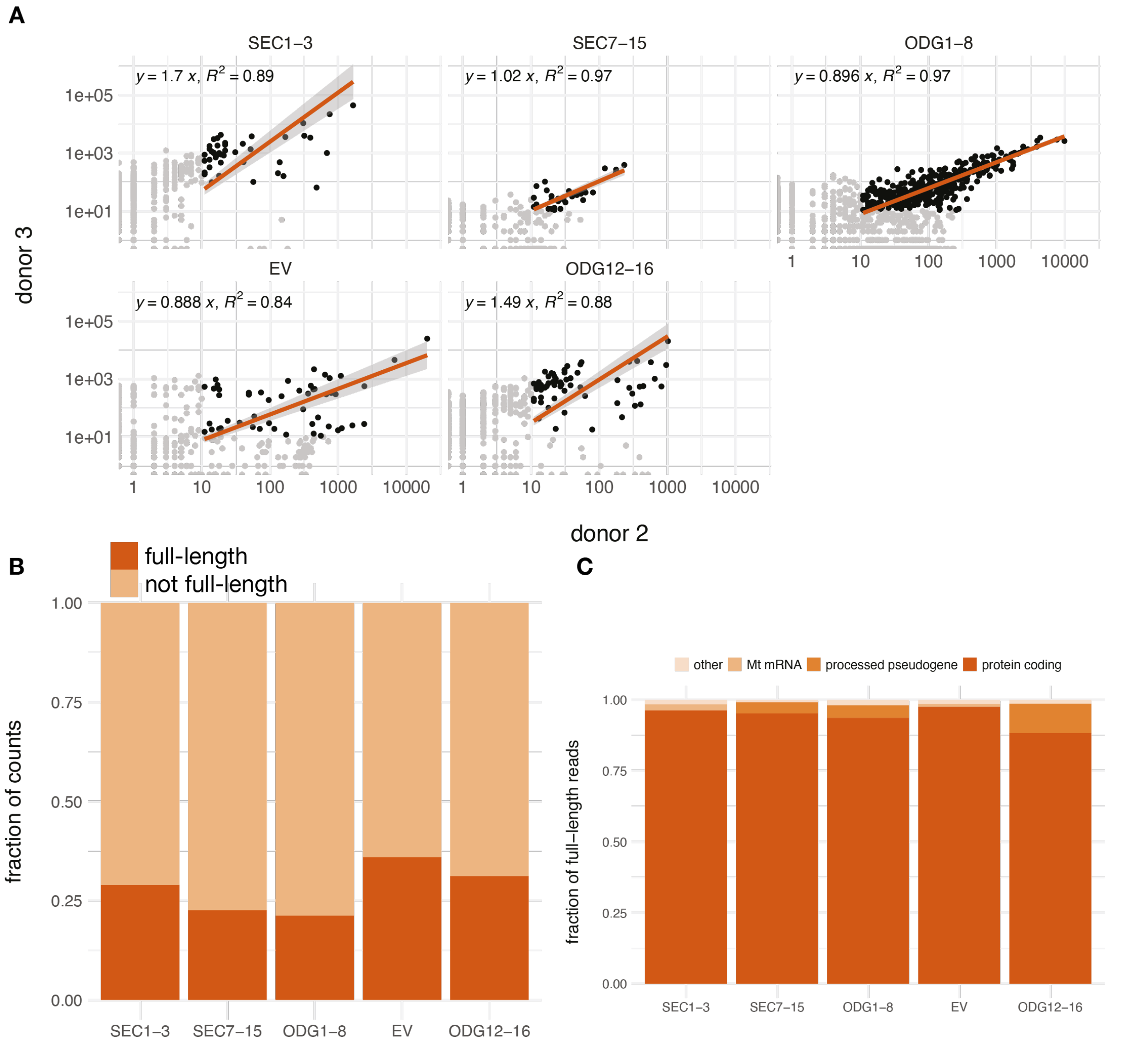


**Supplemental Figure 4: Exploratory analysis of long-read sequencing of PFP fractions.** (A) Correlation plots for biological replicates. For each of the pooled fractions, the x-axis shows the raw counts for donor 2 and the y-axis shows the raw counts for donor 3 (the axes are log10 transformed). Each dot is an isoform. For the correlation calculations, only isoforms with at least 10 counts in each of the biological replicates (colored in black) are used. The equation of the linear model (forced through 0) are shown with accompanying $R^2$ values. (B) The average fraction of full-length reads for each of the fractions. Fractions are calculated by summing and dividing all intact reads by all reads from both samples. (C) Gene biotype distribution for the full-length reads in each of the PFP fractions. The y-axis shows the fraction of the total full-length reads that was assigned to a specific gene biotype. All biotypes that contributed less than 1% of full-length reads is grouped under "other". (D) The length distributions of isoforms with at least one full-length read for each PFP fraction. Boxplots are drawn over the individual dots, outliers from the boxplots are removed (but are represented in the raw data beyond the boxplot lines). For the isoforms with a theoretical length larger than 1.5 kilobases, the gene names are shown.

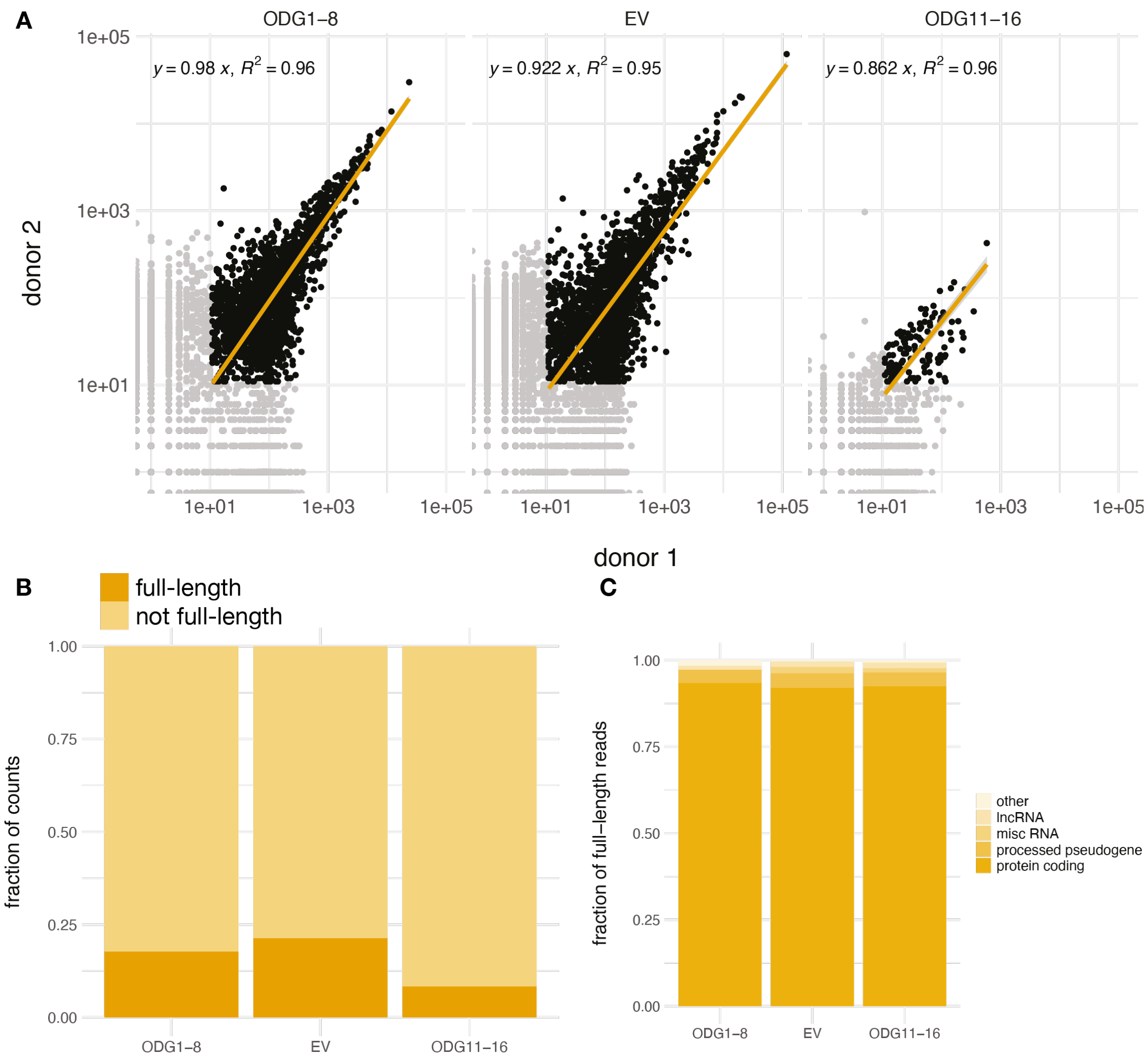


**Supplemental Figure 5: Exploratory analysis of long-read sequencing of urine fractions.** (A) Correlation plots for biological replicates. For each of the pooled fractions, the x-axis shows the raw counts for donor 1 and the y-axis shows the raw counts for donor 2 (the axes are log10 transformed). Each dot is an isoform. For the correlation calculations, only isoforms with at least 10 counts in each of the biological replicates (colored in black) are used. The equation of the linear model (forced through 0) are shown with accompanying $R^2$ values. (B) The average fraction of full-length reads for each of the fractions. Fractions are calculated by summing and dividing all intact reads by all reads from both samples. (C) Gene biotype distribution for the full-length reads in each of the urine fractions. The y-axis shows the fraction of the total full-length reads that was assigned to a specific gene biotype. All biotypes that contributed less than 1% of full-length reads is grouped under "other".
